# PEDIGREE ANALYSIS OF 222 ALMOND GENOTYPES REVEALS TWO WORLD MAINSTREAM BREEDING LINES BASED ON ONLY THREE DIFFERENT CULTIVARS

**DOI:** 10.1101/2020.06.16.154021

**Authors:** Felipe Pérez de los Cobos, Pedro J. Martínez-García, Agustí Romero, Xavier Miarnau, Iban Eduardo, Werner Howad, Federico Dicenta, Rafel Socias i Company, Maria J. Rubio, Thomas M. Gradziel, Michelle Whirthensohn, Henri Duval, Doron Holland, Pere Arús, Francisco J. Vargas, Ignasi Batlle

## Abstract

Loss of genetic variability is a steadily increasing challenge in tree breeding programs due to the repeated use of a reduced number of founder genotypes. High-quality pedigree data of 222 almond [*Prunus dulcis* (Miller) D.A. Webb, syn. *P. amygdalus* (L) Batsch] cultivars and breeding selections were used to study global genetic variability in modern breeding programs from Argentina, Australia, France, Greece, Israel, Italy, Russia, Spain and the USA. Inbreeding coefficients, pairwise relatedness and genetic contribution were calculated for these genotypes. The results reveal two mainstream breeding lines based on three cultivars from two different geographical regions: ‘Tuono’-‘Cristomorto’ (local landraces from Puglia, Italy) and ‘Nonpareil’ (chance seedling selected in California, USA, from French original stock). Direct descendants from ‘Tuono’ or ‘Cristomorto’ number 75 (sharing 30 descendants), while ‘Nonpareil’ has 72 direct descendants. The mean inbreeding coefficient of the analyzed genotypes was 0.036, with 13 genotypes presenting a high inbreeding coefficient, over 0.250. Breeding programs from the USA, France and Spain showed inbreeding coefficients of 0.067, 0.050 and 0.034, respectively. According to their genetic contribution, modern cultivars from Israel, France, the USA, Spain and Australia, trace back to six, five, four, four and two main founding genotypes respectively. Among the group of 65 genotypes carrying the *S_f_* allele for self-compatibility, the mean relatedness coefficient was 0.133, with ‘Tuono’ as the main founding genotype (23.75% of total genetic contribution). Increasing as well as preserving current genetic variability is required in almond breeding programs worldwide to assure genetic gain and continuing breeding progress. Breeding objectives, apart from high and efficient productivity, should include disease resistance and adaptation to climate change. Ultimately, any new commercial almond cultivar has to be economically viable and breeders play a critical role in achieving this goal.

## INTRODUCTION

Almond *Prunus dulcis* is the most economically important temperate tree nut crop worldwide. Due to increasing demand, production areas are expanding into warm and cold climatic regions of both hemispheres. Almond world production (1,258,324 kernel tonnes) is led by the USA (80%), Australia (6%) and Spain (5%) ^1^. The origin of almond within the Amygdalus subgenus, including cultivated almond and its wild relatives such as *P. fenzliana* Fritsh, *P. bucharica* (Korsh.) Fedtsch, *P. kuramica* (Korsh.) Kitam. and *P. triloba* Lindl. ^2–4^ took place approximately 5.88 million years ago ^5^. Almond originated in the arid mountainous regions of Central Asia, where it was first cultivated around 5000 years ago ^6^ and then moved to the Mediterranean region and later to California and the southern hemisphere (South America, Australia and South Africa) ^3^. *P. dulcis* is characterized by its adaptation to Mediterranean climates. Wide cultivation of almond, often under the more severe environments of Central Asia to the Mediterranean region, was possible because of the availability of a highly diverse gene pool, genetic recombination promoted by its self-incompatible mating system and, possibly, by interspecific hybridization and gene introgression involving other members of the Amygdalus subgenus. As a result, almond is an extremely variable species, with a high morphological and physiological diversity. This variability, as measured with biochemical and molecular markers ^7–9^, that has revealed that almond is the most genetically variable of the diploid *Prunus* cultivated species ^10,11^.

In the Mediterranean Region, two thousand years of almond culture have concentrated production to specific areas where well-defined seedling ecotypes and local cultivars have evolved ^2^. By the turn of the 20 th century, most of these almond producing countries had identified locally desirable cultivars that were often seedling selections of unknown origin ^12^. Thus, growers selected cultivars and landraces which represented a rich genetic diversity. Most of these Mediterranean local cultivars have largely disappeared from cultivation in the last 50 years ^13^. Modern almond cultivation is based on a reduced number of cultivars (preferably self-compatible) grafted onto soil adapted clonal rootstocks and under irrigated conditions when possible.

Modern almond breeding started in the 1920’s with the making of controlled crosses and seedling selections to meet changing agronomic and market demands. Currently, there are six active public breeding programs: the USA (UCD-USDA), Spain (CITA, IRTA and CEBAS-CSIC), Australia (University of Adelaide-ABA) and Israel (ARO). Some private breeding programs exist also in the USA. In addition, there were various breeding initiatives in Russia, France, Greece, Italy and Argentina ^13^. Almond breeders have relied mainly on outcrossing and, occasionally, on introgression from other *Prunus* species, for the development of new cultivars. Compared to other fruit tree species, almond breeding had a later start. Initially, in the USA (with limited accessible genetic resources) and later in Russia and Mediterranean region (with more diverse germplasm available) rapid genetic advances occurred. In California, ‘Carmel’ (introduced in 1966), as ‘Nonpareil’ pollinizer, was the first cultivar release with extensive commercial impact. In Russia and the former Soviet Union, several late flowering and frost hardy cultivars were obtained in the 1950’s with ‘Primorskyi’ (date unknown) later used extensively for breeding. In the Mediterranean region, late flowering, productive, well-adaptated and resilient cultivars like ‘Ferragnès’ (1973) or ‘Masbovera’ (1992) were released with great success. The French self-compatible cultivar ‘Lauranne’ (1991) showed a broad environmental adaptation, high production and regular cropping. Although improved cultivars continued to be released, the amount of progress per generation diminishes since parents were continually drawn from the same genepool ^13^. Inbreeding depression expressed as low vigor, reduced flower number and fruit set, increased fruit abortion, lower seed germination and seedling survival, increased leaf and wood abnormalities, and loss of disease resistance have been reported (Grasselly, 1976; Grasselly & Olivier, 1981; Martínez-García, Dicenta, & Ortega, 2012; Socias i Company, 1990). In addition, low self-fruitfulness in self-compatible almond genotypes was suspected to be due to inbreeding ^18^.

Different breeding objectives were developed according to regional commercial and market requirements. Regarding nut shell hardness, two types of almonds are bred (soft-shelled in the USA and Australia mainly) and hard-shelled in most Mediterranean countries. A common aim of these breeding programs was obtaining self-compatible genotypes as most traditional almond cultivars are self-incompatible. Self-compatibility is controlled by a single self-compatibility *S_f_* dominant allele ^19^. During the last 50 years, almond breeding for self-compatibility has mainly used two sources of *S_f_*, local landraces originated in Italy (‘Tuono’ and ‘Genco’) and related species as *P. persica* and *P. webbi* ^20^.

Male parents carrying the *S_f_* allele and sharing the other S-alelle with the female parent are used when breeding for self-compatibility. In addition, crossing heterozygous self-compatible parents in breeding programs has been suggested to obtain homozygous self-compatible genotypes to be used in further breeding ^21^. Such breeding strategies can narrow the genetic variability of crops when they lead to a reduced number of genotypes utilized as parents. Moreover, loss of genetic variability can lead to crop losses due to poor adaptation to biotic and abiotic stresses and inbreeding depression ^22^. Consequently, it’s critical to have accurate knowledge of the relationships among genotypes utilized in breeding programs.

Several almond populations have been analyzed with molecular markers in order to determine genetic variability and relatedness ^9,23–25^. However, these studies were performed with material from limited geographic areas and do not represent the current worldwide status of almond breeding germplasm. Although genomic measures of inbreeding are more accurate than those obtained from pedigree data ^26,27^, pedigree-based analysis is a cost-effective technique to estimate these parameters in breeding populations. Several reports have evaluated inbreeding based on pedigree data in breeding populations of fruit and nut tree crops ^28–31^. In almond, a pedigree analysis of 123 different genotypes from the USA, France, Spain, Israel and Russia was reported ^32^. However, their work was primarily focused on North American genotypes and does not include the many cultivars have subsequently been released worldwide.

This study aims to determine the inbreeding coefficient, pairwise relatedness and genetic contribution of main founding clones, based on pedigree data of 222 almond cultivars and selections released in the last decades in Argentina, Australia, France, Greece, Israel, Italy, Russia, Spain and the USA. We analyzed breeding pedigree at four levels: worldwide, by country with relevant breeding programs, by programs (when different programs exist within a country) and by genotypes carrying the *S_f_* allele for self-compatibility. As far as we know, this is the largest pedigree analysis ever performed in almond.

## MATERIALS AND METHODS

### Pedigree data

Pedigree data of 222 almond genotypes (cultivars and breeding selections) were collected from high-quality sources, including available literature and personal communications from breeders. (Pedigree data and references are present in Supplementary Table 1). In addition, parental relationships for several cultivars were checked using SSRs markers of the IRTA’s almond marker database (unpublished data) and different studies of genetic variability in almond ^9,23–25^. Pedigrees of 170 genotypes of known origin were retrieved (102 cultivars and 68 breeding selections), of which: 59 from Spain, 56 from the USA, 17 from Russia, 11 from Israel, 10 from France, 7 from both Australia and Greece and 2 from both Argentina and Italy.

A pedigree data file was created. Each record in the file contained one cultivar or selection name, the female parent and the male parent, in that order. Once entered, these data were available for inbreeding analyses such as determining the number of times a cultivar appeared in a pedigree as a male or female genitor. Genotypes of known origin were classified into two groups according to self-compatibility (105 self-incompatible and 65 self-compatible).

### Inbreeding coefficient, pairwise relatedness and genetic contribution

The inbreeding coefficient (*F*) is defined as the probability that a pair of alleles at any locus in an individual are identical by descent and it is given by the following formula ^33^:

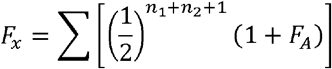

Where *n*_1_ = number of generations from one parent back to the common ancestor, *n*_2_ = number of generations from the other parent back to the common ancestor and *F_A_* = inbreeding coefficient of the common ancestor.

Pairwise relatedness (*r*) or coancestry coefficient, the degree of relationship by descent of two parents, equals the inbreeding coefficient of their prospective progeny.

The genetic contribution (*GC*) of a founder to a cultivar is given by the following formula ^34^:

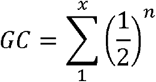

Where *n* = number of generations in a pedigree pathway between the founding clone and the cultivar and *x* = number of pathways between the founding clone and the cultivar. The three parameters were calculated using the SAS INBRED procedure (SAS 9.4 SAS Institute, Cary NC USA).

In summary, the inbreeding coefficient measures the probability that two alleles in a locus are identical by descent and so copies of the same allele from a previous generation. The pairwise relatedness measures the probability that two alleles at any locus are identical by descent (copies of the same allele in a previous generation) between two different individuals. *F* and *r* values range from 0 to 1, with values close to 0 indicating a low degree of inbreeding or relatedness and values close to 1 indicating a high degree of inbreeding or relatedness. The genetic contribution estimates the proportion of genome that comes from the same individual. Thus, a son will have 0.5 genome of either parent and a grandson will have 0.25 genomes of his grandparents.

### Analysis description

To calculate *F*, *r*, and *GC*, parents of unknown origin were assumed to be unrelated and noninbred. The seed parent involved in all open-pollinations was also assumed to be unrelated to the pollen parent. These assumptions, based on the fact that most almond cultivars are obligate outcrossers because of their self-incompatibility, may lead to an underestimation of inbreeding. In the case of genotypes of open pollinated origin (OP); numbers OP1, OP2, OP3, etc. were given in order to be distinguishable for genetic studies. Also, all mutants were considered to have no genetic differences from the original cultivar, thus *GC* = 1. Since the differences between such mutants and the original cultivar are expected to be caused by a few mutations in the DNA, this simplification avoids the overestimation of inbreeding coefficients. Cultivars like ‘Supernova’ and ‘Guara’ were considered as ‘Tuono’ clones ^35,36^. Regarding the different clones of the French paper-shell cultivar ‘Princesse’, used in both the USA and Russian breeding programs, we adopted the approach of Lansari et al. (1994) by analyzing both clones as the same cultivar. Historical reports suggest that the Hatch series ‘Nonpareil’, ‘I.X.L.’ and ‘Ne Plus Ultra’ were seedling selections from an open pollination progeny of the early introduced cultivar ‘Princesse’. This cultivar probably originated from the Languedoc region in France ^37–40^. Because their specific origins remain uncertain, we analyzed these genotypes as non-related, which, however, could lead to an underestimation of inbreeding. Cultivar ‘Mission’ (syn. ‘Texas’ or ‘Texas Prolific’) was similarly selected in Texas about 1891 as a chance seedling of unknown origin ^41^.

Pedigree data were analyzed at four levels: worldwide, by country (Australia, France, Israel, Spain and the USA), by breeding program (when exist different programs within a country: CITA, IRTA, CEBAS-CSIC and, UCD-USDA) and by genotypes carrying the *S_f_* allele for self-compatibility.

## RESULTS

### Founding clones

The entire almond pedigree traces back to 52 founding clones (Supplementary Figure 1). ‘Nonpareil’, ‘Cristomorto’, ‘Mission’ and ‘Tuono’ were the founders with the highest influence on pedigree: 139 of the 170 genotypes of known parentage trace back to one or more of these founding clones (Figure 1). No genotype is derived from all four cultivars, i.e. it does not trace back to the four founding clones.

**Figure 1.**
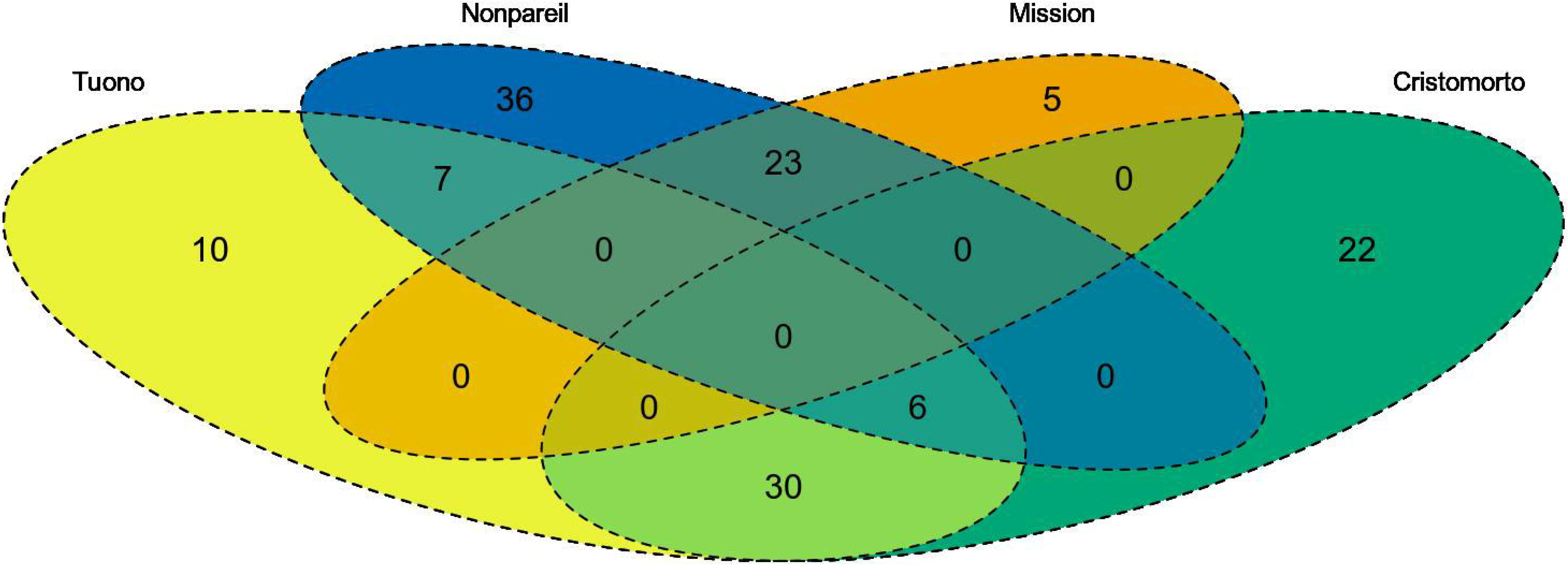
Venn diagram showing the number of descendants shared by ‘Tuono’, ‘Nonpareil’, ‘Mission’ and ‘Cristomorto’.

There are only six genotypes that come from a 3-way shared progeny, all of them tracing back to ‘Tuono’-‘Cristomorto’-‘Nonpareil’. The largest 2-way shared genotype sub in set are ‘Tuono’-‘Cristomorto’ and ‘Nonpareil’-‘Mission’ with 30 and 23 descendants respectively. ‘Mission’ only shares progeny with ‘Nonpareil’ (Figure 1).

Analyzing the results by country, breeding programs from the USA have two main founding clones, ‘Nonpareil’ and ‘Mission’, with 47 and 25 descendants respectively out of 56. These two founders are followed by ‘Eureka’ and ‘Harriott’, with 10 and 7 descendants each. Breeding programs from Spain have three main founding clones, ‘Cristomorto’, ‘Tuono’ and ‘Primorskyi’, with 37, 35 and 24 descendants respectively. Cultivars from the discontinued French program have three main founding clones from two geographical origins, Italy (‘Cristomorto’ and ‘Tuono’) and Southern France (‘Aï’) with nine, four and eight descendants respectively. The Australian program has only two main founding clones, ‘Nonpareil’ and ‘Lauranne’ (‘Ferragnès’ x ‘Tuono’) with six shared derived genotypes out of 7 releases. The Israeli breeding program shows the most balanced pedigree origin with six main founding clones from five different original geographical regions, ‘Marcona’, ‘Greek’, ‘Um ElFahem’, ‘Tuono’, ‘Nonpareil’ and ‘Ferragnès’ out of 11 different genotypes used as parents.

The UCD breeding program has ‘Nonpareil’ as main founding clone with 24 descendants out of 32 used parents. Cultivars ‘Eureka’, ‘Harriott’ and ‘Mission’ have a slight influence in the pedigree with 10, 6 and three descendants respectively. Within Spain, CITA breeding program has Italian ‘Tuono’ as the main founding clone and 7 of 9 released cultivars. The IRTA breeding program shows three main founding clones from two geographical origins, ‘Cristomorto’, ‘Primorskyi’ and ‘Tuono’ with 31, 17 and 14 descendants respectively, with seven released cultivars. The CEBAS-CSIC breeding program has three main founding clones from three different geographical origins, ‘Tuono’, ‘Ferragnès’ and ‘Primorskyi’ with 15, ten and seven descendants respectively and, five released cultivars. The French local cultivar ‘Aï’ has an indirect impact as a grandparent in the three Spanish programs, through the largely used French ‘Ferraduel’ and ‘Ferragnès’ cultivars, are the ancestor of 33 genotypes.

Analyzing the 65 genotypes carrying the *S_f_* allele for self-compatibility, the founding clones which trace back to the origin of this allele are ‘Tuono’, ‘Genco’ and genotypes originated from introgression crosses with *Prunus persica* and *Prunus webbi*. As a source of self-compatibility, the Mediterranean breeding programs have only used Italian local cultivars and the USA programs use ‘Tuono’ and related *Prunus* species.

### Inbreeding coefficients

The mean inbreeding coefficient (*F*) of the 170 genotypes of known parentage analyzed was 0.036 (Supplementary Table 2). Some 36 genotypes presented an *F* > 0, with only 13 over 0.250 (Table 1). The USA accessions ranged from *F* = 0 to 0.375 with ten of the 56 cultivars and advanced breeding lines having *F* > 0.250. The French cultivar ‘Ferralise’ and selection ‘FGFD2’, derived from the same reciprocal cross, being siblings of the same parents, have *F* = 0.250. The Spanish selection ‘A2-198’ from CEBAS-CSIC, showed the highest inbreeding coefficient (*F* = 0.500) as it is a selfing from selection ‘C1328’ and was raised to obtain homozygou *S_f_S_f_* individuals. Considering only the 65 self-compatible genotypes, they had a mean *F* of 0.020, ranging from 0 to 0.150, which is less than the *F* = 0.036 of the 170 grouped genotypes (Supplementary Table 2).

**Table 1.** Genotypes with the highest inbreeding coefficient.

| LineName | FemaleParent | MaleParent | Origin | Country | Inbreeding |
| --- | --- | --- | --- | --- | --- |
| A2-198 | C1328 | C1328 | CEBAS-CSIC | SPAIN | 0.5 |
| Solano | 21-19W | 22-20 | UCD | USA | 0.375 |
| Sonora | 21-19W | 22-20 | UCD | USA | 0.375 |
| Vesta | Nonpareil | Solano | UCD | USA | 0.375 |
| Ferralise | Ferraduel | Ferragnès | INRA | FRANCE | 0.25 |
| FGFD2 | Ferragnès | Ferraduel | INRA | FRANCE | 0.25 |
| 21-19W | Nonpareil | A1-30 | UCD | USA | 0.25 |
| 22-20 | Nonpareil | A1-30 | UCD | USA | 0.25 |
| 6-27 | Nonpareil | Jordanolo | UCD | USA | 0.25 |
| Calif. 24-6 | Eureka | A5-25 | UCD | USA | 0.25 |
| Emerald | Mission | S2 | PRIVATE | USA | 0.25 |
| Profuse | Nonpareil | Jordanolo | PRIVATE | USA | 0.25 |
| Supareil | Nonpareil | Carmel | PRIVATE | USA | 0.25 |

Considering genotypes within each country, the breeding programs showing more inbreeding are those from the USA, France and Spain with 0.067, 0.050 and 0.034 mean *F*, respectively (Supplementary Table 2). The programs from Australia and Israel showed an *F* of 0 as both programs are young and having few breeding generations, respectively. In the case of the Israel program, a broader germplasm was also used. The UCD-USDA breeding program has a mean *F* of 0.083 as their starting germplasm was reduced and has mainly focused on ‘Nonpareil’ types. Broadly, the USA almond breeding activity shows an *F* = 0.067. Within Spain, mean inbreeding was 0.033, the CITA program has *F* = 0, the CEBAS-CSIC program presented an average of 0.024 but only one genotype with *F* > 0 (‘A2-198’). The IRTA program holds 15 genotypes with *F* > 0 and a mean *F* of 0.044 (Supplementary Table 2).

### Genetic contribution

‘Nonpareil’, ‘Tuono’, ‘Cristomorto’ and ‘Mission’ are the founding clones with the highest mean genetic contribution (*GC*; Figure 2). These four cultivars account to 48.6% of the total *GC* worldwide. ‘Nonpareil’ represents 20% of *GC* worldwide, ‘Tuono’ and ‘Cristomorto’ are around 11% *GC* and ‘Mission’ slightly exceeds 5% *GC*. Nevertheless, the mean *GC* of these founding clones within each country is variable. The breeding programs most dependent on these founders include those in Australia and France, where ‘Nonpareil’, ‘Tuono’ and ‘Cristomorto’ represent more than 60% of the total *GC*. Israel is the least dependent country and these founders represented approximately 25% of the total *GC*. Cultivar ‘Nonpareil’ is the most influential founder in the USA and Australia while Spain and France are more based on ‘Tuono’ and ‘Cristomorto’. The cultivar ‘Mission’ was used only in the American programs.

**Figure 2.**
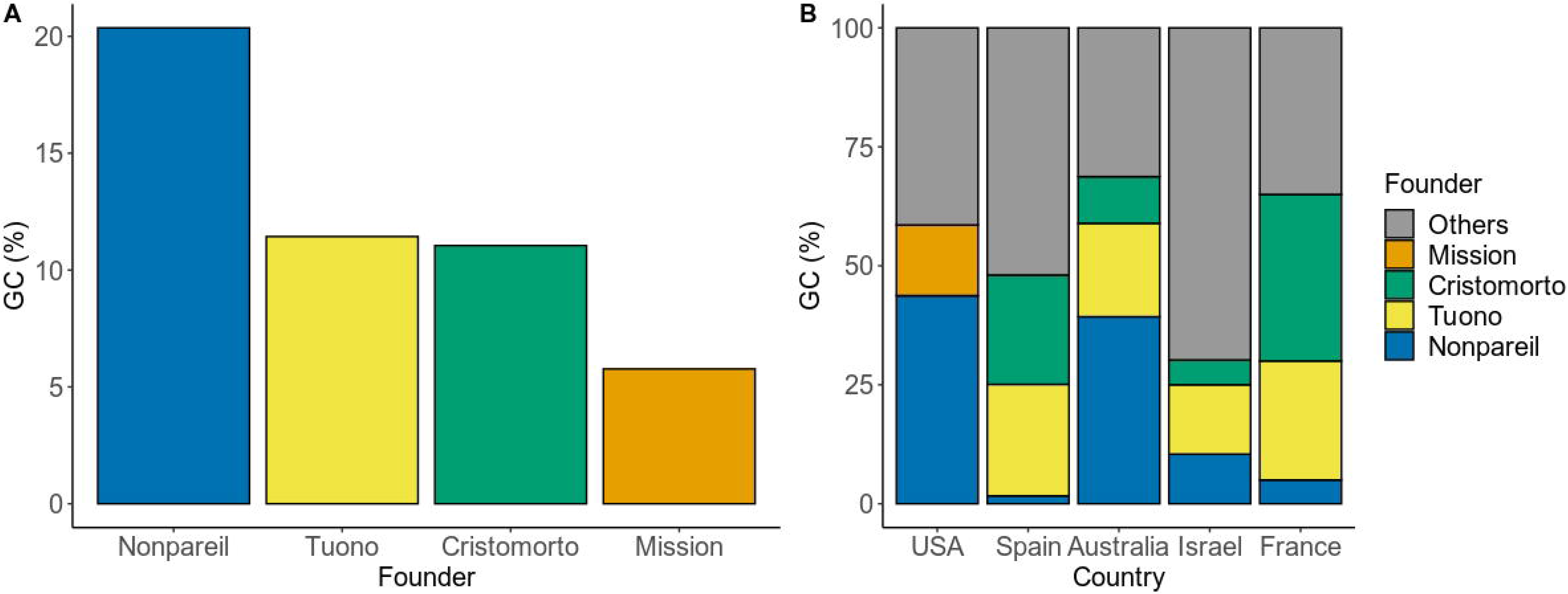
(A) Worldwide mean GC of founders ‘Nonpareil’, ‘Tuono’, ‘Cristomorto’ and ‘Mission’. (B) Mean GC of ‘Nonpareil’, ‘Tuono’, ‘Cristomorto’ and ‘Mission’ by country.

**Table 2** shows the *GC* of the mean founders by country. In the Australian breeding program, only two founders, ‘Nonpareil’ and ‘Lauranne’, represent the 78.57% of the total *GC*. The French breeding program is characterized by the extensive use of three founders ‘Cristomorto’ (*GC* = 35%), ‘Aï’ (*GC* = 30%) and ‘Tuono’ (*GC* = 25%). These cultivars together with ‘Ardèchoise’ and ‘Tardy Nonpareil’ (both *GC* = 5%) account for 100% of the total *GC*. The Israeli breeding program presents six main founders, ‘Greek’ (*GC* = 18.75%), ‘Marcona’ (*GC* = 16.67%), ‘Tuono’ (*GC* = 14.58%), ‘Um ElFahem’ (*GC* = 12.50%) and, ‘Ferragnès’ and ‘Nonpareil’ (both *GC* = 10.42%), which together account for 83.34% of the total gene contribution. The USA breeding programs are largely dependent on ‘Nonpareil’ (*GC* = 43.74%) followed by ‘Mission’ (*GC* = 14.84%), ‘Eureka’ (*GC* = 8.71%), and ‘Harriott’ (*GC* = 5.47%) which all account for 72.75% of the total gene contribution. The cultivars released by the three Spanish breeding programs are based mainly on four founders: ‘Cristomorto’ (*GC* = 23.96%), ‘Tuono’ (*GC* = 22.55%), ‘Primorskyi’ (*GC* = 12.77%) and ‘Aï’ (*GC* = 10.73%), accounting for 70.02% of the total gene contribution.

**Table 2.**
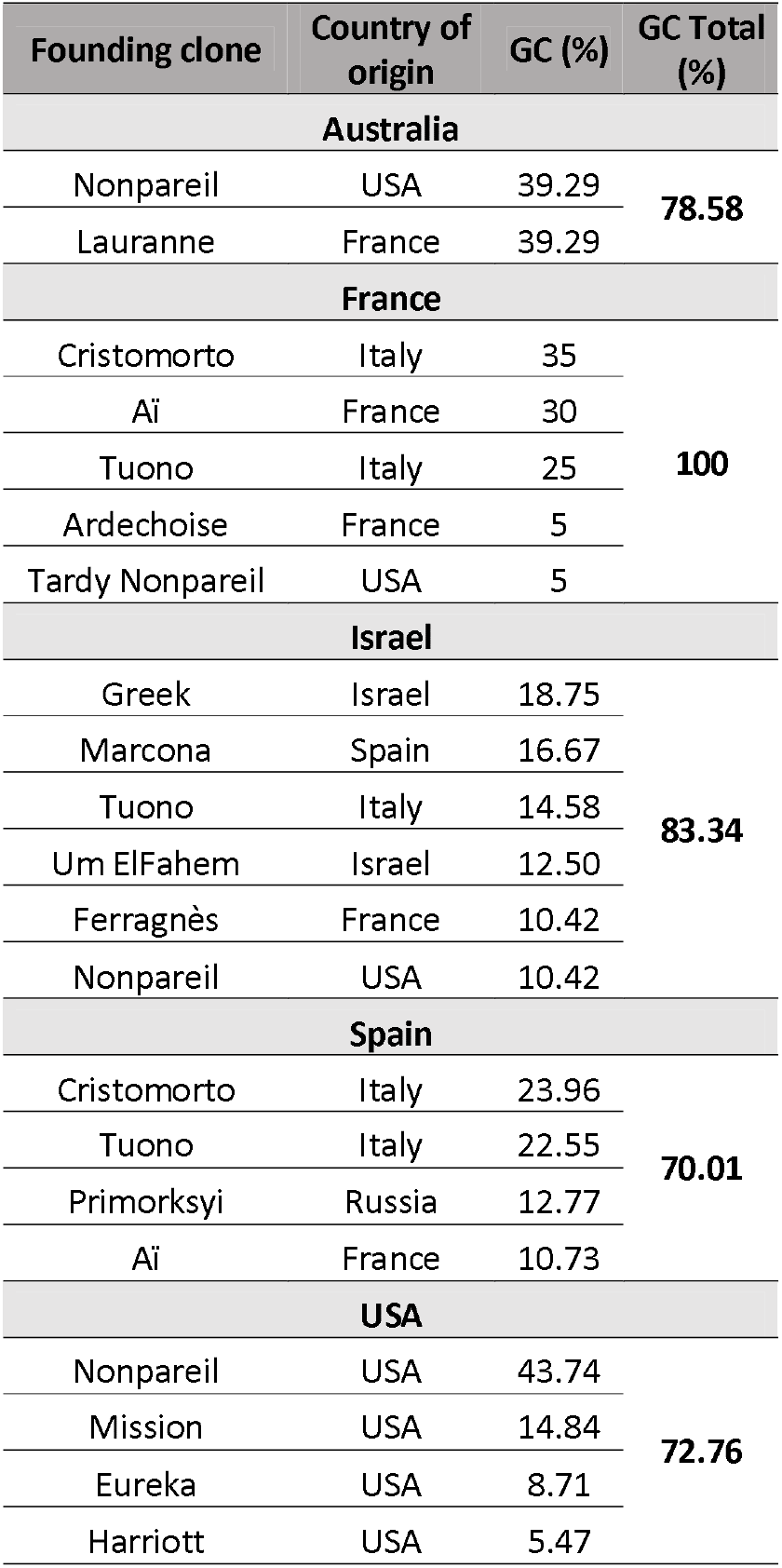
Genetic contribution (*GC*) of mean founding clones by country.

The UCD-USDA breeding program has the same founders than the overall American programs, ‘Nonpareil’ (*GC*=42.94%), ‘Eureka’ (*GC*=15.23%), ‘Harriott’ (*GC*=8.79%) and ‘Mission’ (*GC*=5.65%). In this case, ‘Mission’ influence is reduced. Differences appeared in the use of founding cultivars between Spanish breeding programs. Thus, the CITA program is mainly based on four cultivars ‘Tuono’ (*GC*=35%), and ‘Belle d’Aurons’, ‘Bertina’ and ‘Genco’ (*GC*=10% each). These cultivars are accounting for 65% of the total gene contribution. This program is including a local Spanish late flowering cultivar ‘Bertina’ as founder and also used other late flowering local selection AS-1 which was deduced to be the parent of ‘Blanquerna’ (Fernández i Martí et al. 2015). The CEBAS-CSIC program is based also on four founders, ‘Tuono’ (*GC*=30.36%), ‘Ferragnès’ (‘Cristomorto’ x ‘Aï’) (*GC*=19.05%), ‘Genco’ (*GC*=11.31%) and ‘Primorskyi’ (*GC*=10.12%). The IRTA program is based on four founding clones too: ‘Cristomorto’ (*GC*=35%), ‘Primorskyi’ (*GC*=18.07%), ‘Tuono’ (*GC*=15.26%) and ‘Aï’ (*GC*=13.26%) through ‘Ferragnès’. Some 23 cultivars and selections from IRTA program are only derived from these four founders (Supplementary Table 3). The self-compatible Italian cultivar ‘Tuono’ was the most commonly used genitor by the three programs. Within the 65 genotypes bred carrying the *S_f_* allele, ‘Tuono’ represents on average the 23.75% of the total *GC*, which means being donor in 73.80% of the obtentions (48), with variable contribution according to programs (Supplementary Table 3).

### Pairwise relatedness

Pairwise relatedness (*r*) between all cultivars and breeding selections are showed in Supplementary Table 4. Cultivars with the highest mean *r* worldwide are present in **Table 3**. The genotype with the highest mean *r* is ‘Tardy Nonpareil’ followed by ‘Nonpareil’ and the rest of its mutants (‘Jeffries’ and ‘Kern Royal’). Although ‘Tardy Nonpareil’ appears to be a late-flowering budsport mutation of Nonpareil, its use in both French and Spanish breeding programs explains it higher *r* value than ‘Nonpareil’ and its other mutants. Australian genotypes, which originated from the cross ‘Nonpareil’ by ‘Lauranne’, follow. These genotypes are first generation of ‘Nonpareil’, second generation of ‘Tuono’ and third generation of ‘Cristomorto’, causing their high degree of relatedness with the rest of genotypes.

**Table 3.** Genotypes with the highest mean relatedness (*r*).

| Genotype | Mean $r$ |
| --- | --- |
| Tardy Nonpareil | 0.568 |
| Nonpareil | 0.152 |
| Jeffries | 0.152 |
| Kern Royal | 0.152 |
| Vesta | 0.142 |
| A97001-1bT4 | 0.134 |
| Capella | 0.133 |
| Carina | 0.133 |
| Mira | 0.133 |
| Maxima | 0.133 |

**Table 4** shows the mean *r* among breeding programs by country. Programs from Australia and France have the highest mean *r* values (0.310 and 0.357 respectively), due to the repeated use of relatively few founders, (two and five respectively). In contrast, Israel shows the lowest mean *r* as six founders were reported. Comparing relatedness results between countries, Spain and the USA breeding programs are the less related, as Spanish programs are based on ‘Tuono’-‘Cristomorto’ and the USA programs are based on ‘Nonpareil’-‘Mission’. The most related breeding programs are those of France and Spain and also, Australia and France, showing the strong influence of the French germplasm in both countries.

**Table 4.** Mean of pairwise relatedness (*r*) among breeding programs from five different countries.

|  | Australia | France | Israel | Spain | USA |
| --- | --- | --- | --- | --- | --- |
| Australia | **0.310** | 0.187 | 0.089 | 0.113 | 0.172 |
| France | - | **0.357** | 0.070 | 0.195 | 0.023 |
| Israel | - | - | **0.134** | 0.049 | 0.050 |
| Spain | - | - | - | **0.163** | 0.009 |
| USA | - | - | - | - | **0.235** |

In the Australian breeding program, the selection ‘A97001-1BT4’ has the highest mean *r* with a value of 0.417. ‘Rhea’ is not related with the rest of the genotypes so its mean *r* is zero. The rest of the genotypes have a mean *r* values between 0.25 and 0.375 showing a highest degree of relationship.

In the French breeding program, ‘Ferralise’ has the highest mean *r* (0.500), this cultivar originating from the cross ‘Ferragnès’ by ‘Ferraduel’, which are two sibling cultivars. ‘Ferrastar’ and ‘R1000’ have the lowest mean *r* values, 0.167 and 0.111 respectively. The rest of French genotypes have a mean *r* values over 0.300.

The Israeli program bred the least related genotypes, as none of them has a mean *r* of over 0.225. The highest *r* value observed between the ten cultivars released was 0.500 between two pairs: ‘Dagan’-‘Gilad’ and ‘Fergil’-‘Gilad’. Selection ‘54’ showed *r* of 0.500 with ‘Kochba’ and 0.250 with ‘Kogil-Pat’, ‘Samish’ and ‘Solo’. Figure 3 shows the relationships of genotypes from Israel and France.

**Figure 3.**
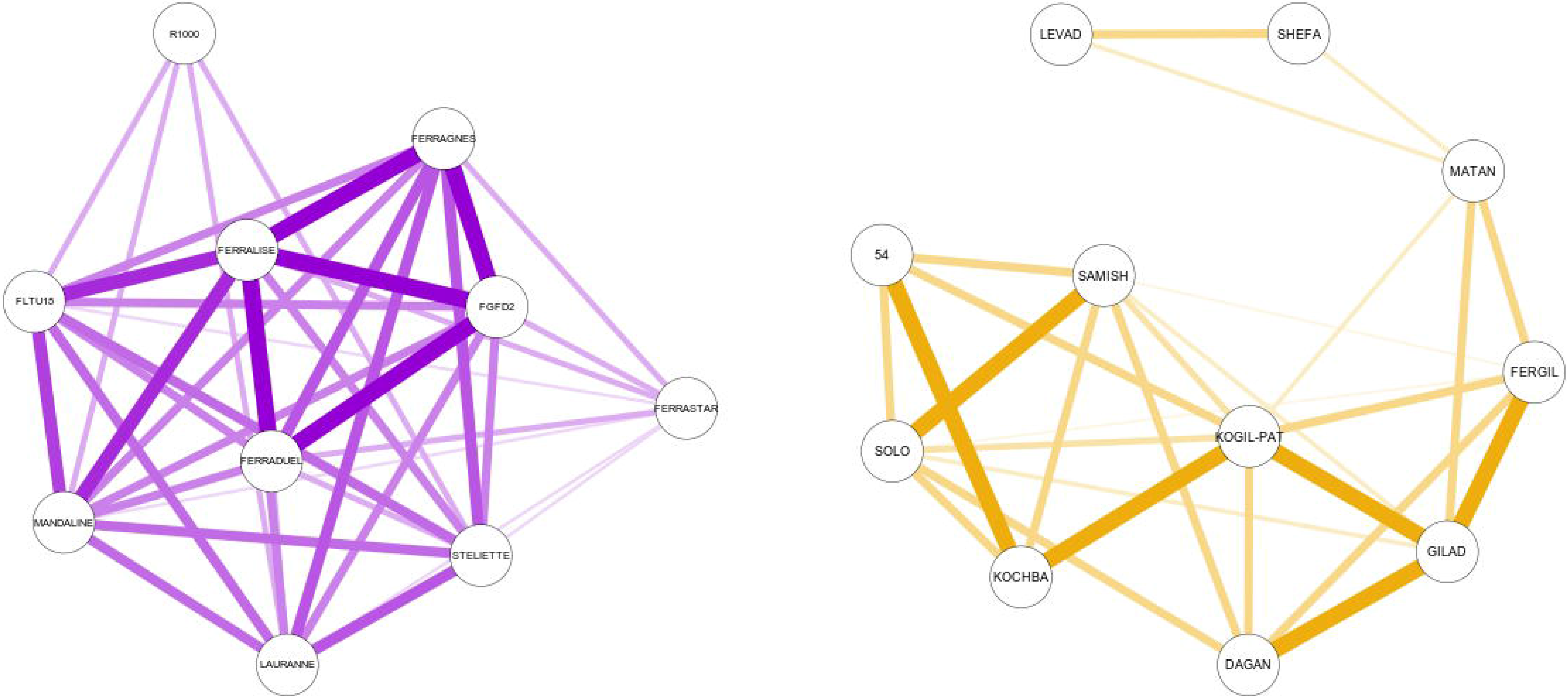
Relationship matrix of genotypes from France (left) and Israel (right).

Within the Spanish breeding programs, the highest *r* among released cultivars was 0.500 between cultivars ‘Antoñeta’-‘Marta’ and ‘Makako’-‘Penta’ as both couples are siblings derived from the same cross (‘Ferragnès’ x ‘Tuono’). The cultivar pairs, ‘Makako’-‘Tardona’ and ‘Penta’-‘Tardona’ have an *r* = 0.313 as both crosses have a common parent in selection ‘S5133’. The CEBAS-CSIC’s selections ‘A2-192’ and ‘C1328’ have the highest *r* with a value of 1 (‘A2-198’ was originated from a self-pollination of ‘C1328’). In the CEBAS-CSIC program, ‘D01-462’ has the highest mean *r* (0.273). The genotypes with a higher mean *r* in the CITA breeding program are ‘Guara’ and ‘Felisia’ with values of 0.278 and 0.250 respectively. The remaining CITA genotypes present a mean *r* values of under 0.200. At IRTA breeding program, the highest *r* value among released cultivars was 0.563, between two cultivar pairs ‘Anxaneta’-‘Tarragonès’ and ‘Glorieta’-‘Marinada’, showing parent-offspring relation. Among IRTA’s selections, ‘29-47’ and ‘35-164’, showed the highest relationship with an *r* of 0.719. The selection ‘29-47’ presents the highest mean *r* (0.350). The selection ‘21-321’, is the most related to the rest of the genotypes from Spain with a mean *r* of 0.244. The rest of IRTA’s genotypes have mean *r* values over 0.130 (Supplementary Table 4).

In the USA breeding programs, all ‘Nonpareil’ mutations have a mean *r* values over 0.400. ‘Independence’ and ‘Bell’ gave a mean *r* equal to 0. The rest of North American genotypes showed a high degree of relatedness between them. Two combinations (‘Solano’-‘Vesta’ and ‘Sonora’-‘Vesta’) had *r* = 1, being ‘Sonora’-‘Vesta’ *r* =0.875. In analyzing the highest *r* values among selections and cultivars, four combinations have an *r* = 1 (‘21-19W’-‘Solano’, ‘22-20’-‘Solano’, ‘21-19W’-‘Sonora’, ‘22-30’-‘Sonora’). In addition, two other pairs: ‘21-19W’-‘Vesta’ and ‘22-20’-‘Vesta’ have an *r* of 0.875 (Supplementary Table 4). Within the UCD breeding program, ‘Vesta’, ‘Sonora’ and ‘Solano’ have a mean *r* over 0.400.

Among the group of 65 genotypes carrying the *S_f_* allele, the mean *r* was 0.133. Grouping the genotypes by origin of the *S_f_* allele source (‘Tuono’, ‘Genco’ and other *Prunus spp*) the mean *r* were 0.219, 0.333 and 0.173 respectively (Supplementary Table 4 and Figure 4). Figure 4 shows the main self-compatibility sources used when breeding for this character with ‘Tuono’, ‘Genco’ and other *Prunus* species involved in 48, 4 and 13 genotypes respectively magnifying the importance of these two Italian cultivars^16,20,42^.

**Figure 4.**
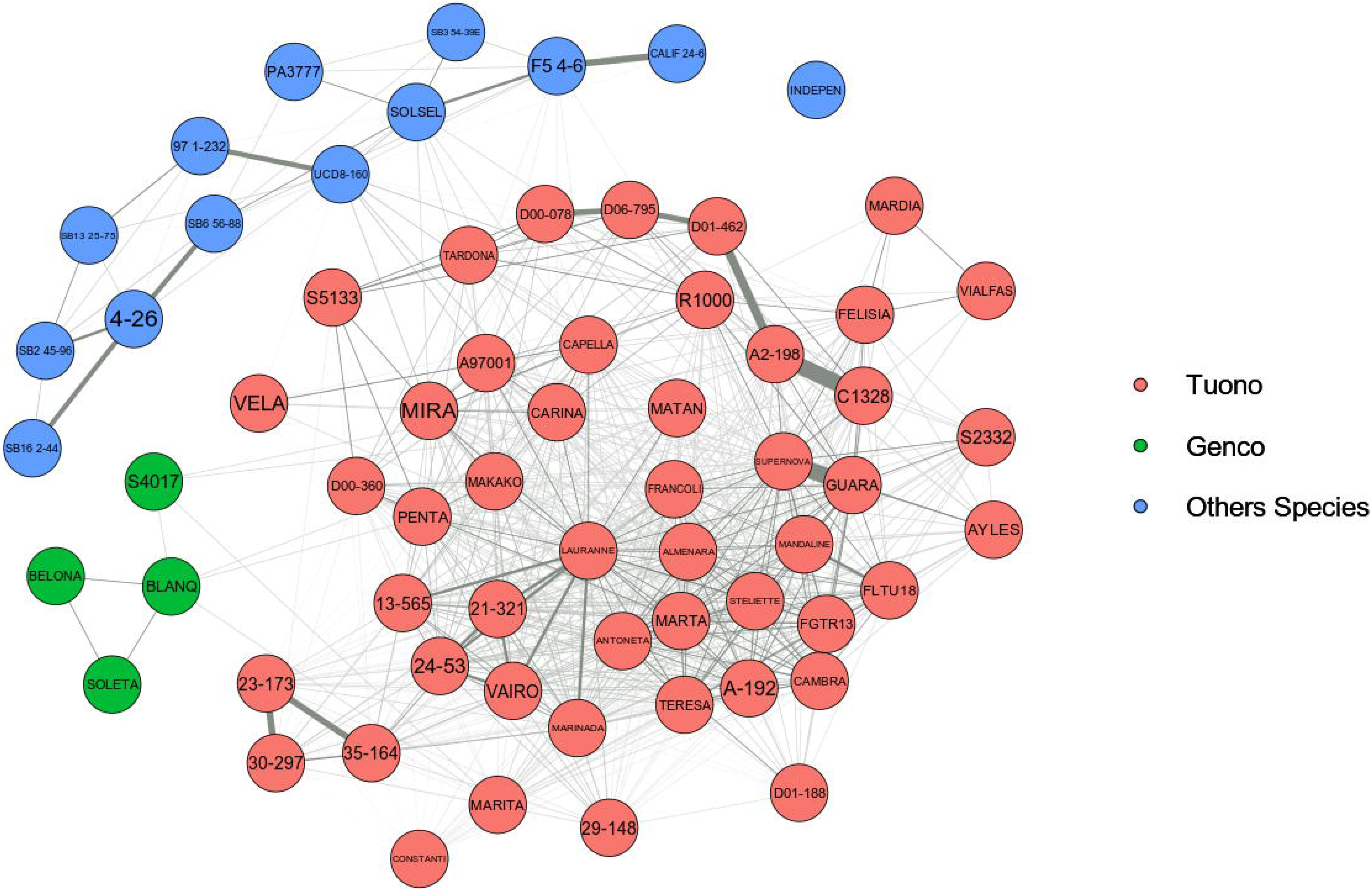
Relationship matrix of the 65 self-compatible genotypes carrying the S_f_ allele and its origin. Some names are shortened.

## DISCUSSION

### Two mainstream breeding lines based on three different cultivars

This genetic study of almond breeding programs worldwide over the last 50 years and based on high-quality pedigree data, demonstrates that the most influential cultivars were ‘Nonpareil’, ‘Tuono’, ‘Cristomorto’ and ‘Mission’. ‘Nonpareil’ had the most influence as it has been extensively used in USA and Australian programs as soft-shelled nuts are bred. This reference cultivar and, in some cases, through its late blooming mutant ‘Tardy Nonpareil’, was present in all the breeding programs studied. In addition, the self-compatible ‘Tuono’ and the late blooming ‘Cristomorto’ were used extensively in the pedigrees analyzed, mainly in the Mediterranean programs, as hard-shelled nuts are bred. ‘Mission’, however, showed a considerable importance worldwide, though partial analysis demonstrated that it was mainly influential in private American programs. These founders had a different relative use depending on country. Thus, we can establish two main breeding lines based on the use of these three different founders: the European programs based primarily on ‘Tuono’ and ‘Cristomorto’ (hard-shell), and the North American-Australian programs based on ‘Nonpareil’ (soft shell). The French and Spanish breeding programs are based directly on ‘Tuono’ and ‘Cristomorto’. In the French INRA program, the Italian cultivars ‘Tuono’ and ‘Cristomorto’ account for 60% of total *GC* and are present in all ten cultivars and selections evaluated. Also, the local French late-flowering and Monilinia resistant cultivar ‘Aï’ was a parent to both ‘Ferragnès’ and ‘Ferraduel’. In the three Spanish breeding programs, the importance of ‘Tuono’ and ‘Cristomorto’ cultivars is very high, accounting to 46.41% of total *GC* and are present in the pedigrees of 52 out of 58 cultivars and breeding selections. In the North American breeding programs, ‘Nonpareil’ accounts for 43.74% of the total *GC* and is present in 47 out of 56 cultivars and breeding selections released. In Australia, ‘Nonpareil’ accounts for 39.29% of the total *GC* and is present in 6 out of 7 cultivars and breeding selections. Even in other countries with non-continuous breeding initiatives, such as Russia, Greece or Argentina, the use of ‘Nonpareil’ as a founder is common. Israel is the only country where these cultivars have a relatively low influence. This may be due to the extreme Israeli climatic conditions, forcing breeders to use locally-adapted selections as parents. In Spain, the use of locally-adapted cultivars such as ‘Bertina’ at CITA as a donor for *Polystigma ochraceum (Wahlenb.) Sacc*. resistance and ‘Marcona’ at IRTA was successful but used only to a limited extent. Other examples of secondary founders include ‘Primorskyi’, used regularly as late-blooming and Fusicoccum resistance donor in two of the Spanish breeding programs (IRTA and CEBAS-CSIC), ‘Eureka’ and ‘Harriott’ in the North American breeding programs and ‘Lauranne’, which achieves a similar importance as ‘Nonpareil’ in the Australian program breeding lineage.

### Loss of genetic variability at breeding and production level

The repeated use of a limited number of parents (‘Nonpareil’, ‘Tuono’ and ‘Cristomorto’) and their related genotypes results in a loss of variability and an associated increase of inbreeding, as we have shown for almond globally. Similar situations have been reported for various rosaceous species including peach^43,44^, strawberry^34^, Japanese plum^45^, apple^46^ and sweet cherry^30^. Comparing our results on almond inbreeding with other *Prunus* species, the mean inbreeding coefficient worldwide of all genotypes (*F*=0.036) is lower than that of Japanese plum ^45^ and apple ^46^ and several orders of magnitude lower than those calculated for peach ^43^ and cherry ^30^.

Among the group of the 65 genotypes carrying the *S_f_* allele for self-compatibility, the mean relatedness coefficient was 0.133. In cherry self-compatible selections, coefficients of coancestry ranged from 0.102 to 0.256 ^30^ and thus were of similar magnitude. In Western Europe, breeders have extensively used the Italian cultivar ‘Tuono’ as a source self-compatibility and more recently it has become important in Israel as well as Australia through the cross ‘Lauranne’ (‘Ferragnès’ x ‘Tuono’). It is interesting to note that this original INRA selection produced the commercially important cultivar ‘Steliette’ and was later successfully utilized in two of the Spanish breeding programs resulting in three self-compatible cultivars ‘Cambra’ at CITA, and ‘Antoñeta’ and ‘Marta’ at CEBAS-CSIC. In addition, in the USA, breeders are using ‘Guara’ (syn ‘Tuono’) as *S_f_* donor. A similar case occurred in sweet cherry with the cultivar ‘Napoleon’ as it was the most frequently utilized parent for self-compatible selections in North America ^30^.

A lack of diverse germplasm may, however, limit continued progress in almond breeding programs. This genetic limitation is of particular concern in the main producing countries. Thus, Californian and Australian production rely mainly on ‘Nonpareil’ and related cultivars^47,48^, while in Spain, some new Spanish cultivars like ‘Vairo’ and ‘Penta’, derived from second generation of ‘Tuono’ and ‘Cristomorto’, as well as ‘Belona’ and ‘Soleta’, derived from second generation of ‘Genco’, are replacing traditional cultivars in new orchards. Only in some regions of Central Asia, Middle East and North Africa, local and well adapted traditional selections still playing an important role in commercial production ^25,49–51^.

### Large-scale genomic data are needed

Pedigree analysis is a cost-effective and well-established way to monitoring inbreeding and relatedness among controlled breeding populations. However, large-scale genomic analysis may outperform analysis ^26,27^. This kind of genome based pedigree analysis has already been performed in apple ^52^. The recent publication of two almond reference genomes ^5,53^ and the increasing availability of quality genomic data opens opportunities to complement this type of study to obtain more complete and accurate pedigrees based on DNA variability.

Also, as commented above, many founders of this pedigree analysis could have the same origin and there are several open pollinated crosses where the male parent information is missing. The use of large-scale genomic data would provide most valuable information in this respect, expanding the almond pedigree beyond breeding records.

### Future breeding lines

There is a need to broaden the genetic base when breeding new almond cultivars. Despite the availability of a large number of modern cultivars and selections from breeding programs worldwide, the size of the genetic resources currently used by breeders is shrinking and, in the course of future genetic improvement, may become insufficient. For continued improvement of commercial traits in almond, genetic gain should be maximized and loss of diversity minimized. Possible methods to accomplish this include using more diverse parents when crossing, using markers to maximize genetic distance during planning, using other sources of self-compatibility, disease resistance and environmental adaptation. In peach, where this drawback is even larger, different strategies based on molecular markers, including marker assisted introgression, have been developed to efficiently introduce useful variability from related species ^54^. In almond, this approach can also be applied using other compatible species of the *Amygdalus* subgenus.

Whereas nut quality parameters are of ultimate importance for industry acceptance, selection for good kernel does not entirely address production and commercial needs. Improved disease resistance, tree architecture, bearing habit, pruning and mechanical harvest remain important traits. Germplasm with the potential of alleviating these problems requires characterization and exploitation and future almond breeding need to expand their germplasm resources. According to Zohary & Hopf (2000), the climatic requirements of fruit and nut tree cultivars closely resemble those of their wild relatives and domestication has not been expanded significantly beyond them. In the context of global warming leading to harsher production environments, new germplasm options need to be characterized, maintained and developed for breeding. Thus, the recovery and introgression of regionally adapted traits from Central Asia, Near and Far East and North Africa represent a promising strategy to develop almonds better adapted to hot climates and limited water availability. Similarly, the use of related *Prunus* species represents a longer-term strategy since several backcrossing generations will be required for successful introgression ^42^. Marker-based approaches such as MAI ^54^ would allow improved selection efficiency and so reduced time. Consequently, future progress in almond breeding will depend on the long-term introgression of novel gene combinations into locally adapted genetic backgrounds.

## CONCLUSIONS

This study of breeding pedigrees reviews and summarizes the progress made in almond breeding over the last 50 years. Results identify two important limitations: the narrowness of modern breeding germplasm utilized and the limited number of generations developed. Substantial improvement in the performance of the new cultivars over traditional cultivars has been achieved. However, results show how two main breeding lineages, based on only three cultivars (‘Cristomorto’, ‘Nonpareil’ and ‘Tuono’) have dominated modern breeding worldwide. This genetic bottleneck risks increased inbreeding depression and reduced genetic variability. Thus, in spite of the high level of available species diversity, caution must be exercised in future almond breeding to avoid inbreeding through the recurrent use of related parents, and to diversify the sources of self-compatibility, which are presently dominated by ‘Tuono’. Broadening the genetic resources available to breeders is an urgent need. Additional analyses based on genomic data are needed to more accurately determine the levels of inbreeding and the loss of genetic variability among almond breeding programs worldwide.

## Acknowledgements

This research was supported in part by grants from the Ministry of Economy and Competitiveness (MINECO/FEDER Projects RTA 2017-00084-00-00 and CERCA Program Generalitat of Catalonia. FPC acknowledges a grant from the Spanish MINECO.

## Author Contributions

F.P.C. and I.B. conceived and designed the study. F.P.C. performed the analysis. F.P.C., P.M., A.R., X.M., I.E., W.H., F.D., R.S., M.R., T.G., M.W., H.D., D.H., P.A., F.V., I.B. wrote the paper. All authors approved the final version of the manuscript.

## Conflict of interest

Authors declare no conflict of interest.

## Notes

### Competing Interest Statement

The authors have declared no competing interest.

