## Supplementary figures and images for "PEDIGREE ANALYSIS OF 222 ALMOND GENOTYPES REVEALS TWO WORLD MAINSTREAM BREEDING LINES BASED ON ONLY THREE DIFFERENT CULTIVARS"

### Supplementary Figure 1

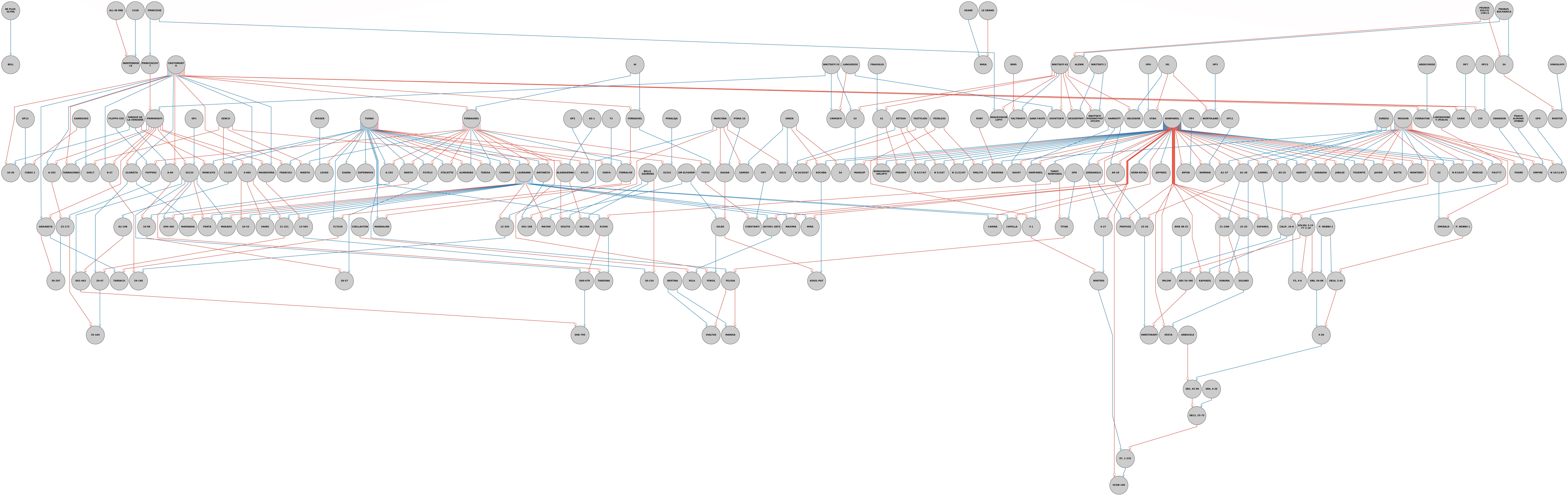
